# OzWheat: a genome-to-phenome platform to resolve complex traits for wheat pre-breeding and research

**DOI:** 10.1101/2024.08.11.603522

**Authors:** Jessica E. Hyles, Howard A. Eagles, Kerrie Ramm, Bjorg Sherman, Andrew Gock, Sandra Stops, Tanya Phongkham, Emmett Leyne, Tina Rathjen, Radoslaw Suchecki, Lauren Stevens, Louise Ord, Nick S. Fradgley, Meredith D. McNeil, Annelie Marquardt, Samuel C. Andrew, Kerrie Forrest, Russell F. Eastwood, Adam Norman, Annette Tredrea, Richard Trethowan, Ben Trevaskis, Shannon K. Dillon

## Abstract

For over a century, Australian wheat breeders have successfully adapted wheat to a broad range of climatic conditions and crop management practices. The OzWheat genome-to-phenome (G2P) platform was established to capture this breeding history and explore traits, genes, and their interactions with the environment to enable ongoing research and deliver targets for wheat improvement. A panel of 285 cultivars and landraces were chosen through knowledge of breeding pedigrees to represent both global diversity and the historic flow of genetic variation over more than 100 years of selective breeding in Australia. Genetic characterisation of the panel included identification of genome-wide sequence variants and gene expression profiling across environments. Important traits for adaptation (flowering time and plant height) were assayed in controlled environments and at multiple field sites and years, with genome-wide association analyses (GWAS) using linear mixed models detecting both known and novel loci. Here, we report establishment of the OzWheat G2P platform as a powerful tool to integrate wheat genomes and phenomes and demonstrate its use to identify candidate genes and understand gene by environment interactions. This provides the wheat research and breeding community a new resource to support future cultivar development.

## Introduction

Wheat is an important food crop worldwide, with global production forecast at 787 million tonnes in 2023/24, representing 28% of total cereal production (FAO 2023). To meet the needs of a growing world population, it is imperative that wheat production is increased and a global research effort to improve wheat yield in changing climates is underway (see Bentley et al. 2022, Fischer et al. 2014). With this challenge in-mind, wheat pre-breeding research has benefitted from extensive development, sharing and deployment of wheat genomic resources to characterise traits which underpin crop performance and identify target genes for crop improvement (Krasileva et al. 2017, Walkowiak et al. 2020, Rogers et al. 2024). Traditional pre-breeding research has typically involved time consuming and labour-intensive approaches such as map-based cloning in bi-parental crosses, development of near-isogenic lines or proof-of-function analysis via transgenesis (Borrill et al. 2018). Although fundamental to defining gene function, these types of analyses in a limited number of genetic backgrounds have not always provided an accurate understanding of gene effects for complex traits. That is, they have not always captured the genetic architecture of polygenic traits nor how genes interact with the environment. Genome-wide association analysis (GWAS) aims to overcome such limitations by surveying a broader genetic base and applying whole-of-genome scale analyses for the simultaneous identification of large effect loci (major genes) together with minor-effect or additive genetic loci (Rafalski 2010).

For GWAS to identify genetic variation which will be relevant to germplasm in a breeding program, it is important to carefully choose the genetic diversity which underpins the platform. Over the course of breeding, populations are developed through crossing and selection with the highest performing, best adapted lines becoming released cultivars. Germplasm sharing between breeders and researchers and frequent intercrossing, backcrossing and selfing to fix lines for release, means that genetic loci are recombined, while recurrent selection and backcrossing maintains favourable allelic combinations. Such germplasm represents a valuable source of diversity for GWAS as it provides opportunity for high-resolution marker-trait associations (Yu et al. 2006). In addition, utilising germplasm which represents the ancestry of current cultivars is potentially a way to validate the phenotypic effects of alleles which have been inherited through a breeding program over time.

Wheat breeding in Australia began over a century ago, when pioneer breeder William Farrer found that colonial wheats were not well-adapted to local growing conditions (Evans, 1980). To create earlier-maturing wheat which flowered at the optimum time for the Australian environment, Farrer made crosses between Fife (hard grain Canadian wheat) and Indian selections which were adapted to high temperatures (Guthrie 1922). By combining favourable quality attributes and high yield potential, Farrer produced a plethora of cultivars which feature in the ancestry of many modern wheats today. Most notably, Federation wheat which was released in 1901 remained the most widely grown cultivar for more than 20 years (Macindoe and Brown 1968).

The enduring success of Farrer wheats can largely be attributed to the development of germplasm adapted to specific growing environments. Today, it is just as important to develop adapted wheat, as crops are cultivated across a broad geographic range, in different farming systems, and in changing climates. Major genes important for adaptation (and therefore yield) include those that affect flowering behaviour (phenology) and plant architecture, and many important loci which affect these traits have been identified including *REDUCED HEIGHT1* (*RHT1), VERNALISATION1 (VRN1), PHOTOPERIOD1 (PPD1)* and *EARLINESS PER SE (EPS)* (Peng et al. 1999, Trevaskis et al. 2003, Diaz et al. 2012, Gawronski and Schnurbusch 2012). Extensive studies in Australia have highlighted the impact of allelic variation of such loci for adaptation (Eagles et al. 2009; Cane et al. 2013; Eagles et al. 2014) although it is apparent that these major genes do not fully explain phenological development in different environments (Bloomfield et al. 2018).

Designing GWAS experiments to allow detection of genetic, environment and their interaction (G×E) effects will therefore be essential to informing our understanding of regulatory mechanisms underlying wheat traits and the reliability of gene targets for breeding. Integrating association analysis across multiple carefully selected environments is becoming a standard approach to address this need supported by a broad set of statistical approaches to partition and detect significant effects (see Tibbs Cortes et al. 2021). The inclusion of other ‘omic data types with GWAS analysis, such the transcriptome or proteome, which are a direct function of G, E and G×E provide an additional avenue for identification of genes which respond to environmental cues and vary in expression level across individuals (Wu et al. 2022, Han et al. 2022, Dillon et al. 2024). This can be achieved by including these ‘omic variables in the association analysis, or likewise through post-GWAS analysis to bolster confidence in identified associations with additional lines of evidence.

This study aimed to develop a Genome-to-Phenome (G2P) platform that provides a new dimension to GWAS, through incorporation of multi-environment assessments and validation of gene associations with gene expression patterns. The addition of gene expression data aims to provide additional insight into the regulatory mechanisms underpinning yield component traits. Phenology and plant height were chosen as exemplar traits to demonstrate the G2P approach using multi-environment, whole-transcriptome variation of a pedigree-informed diversity panel. To handle the extensive datasets created we developed and share an online interface to allow users to visualise and interact with the data by exploring sequence variants, haplotypes and cross-environment gene expression. The OzWheat germplasm and resource connects advances in genomics, transcriptomics and phenomics, providing a G2P platform for the wheat research community to deliver outcomes for breeding.

## Materials and Methods

### Genetic material

The OzWheat Panel consists of landraces, historic releases and modern cultivars chosen to include founders, key introductions and important parents in Australian wheat breeding. The year and region of release in Australia was also considered to ensure a broad range of adaptation and an accurate representation of the flow of alleles through time. Finally, a small number of lines outside the Australian pedigree but with interesting or important agronomic traits or genetic diversity were selected. This selection ensures relevance of the panel to modern Australian breeding programs and growing conditions. In total, 285 cultivars and unreleased breeding lines were sourced directly from breeders, and the Australian Winter Cereals Collection (AWCC, Tamworth, NSW Department of Primary Industries) and Australian Grains Genebank (AGG, https://agriculture.vic.gov.au/crops-and-horticulture/the-australian-grains-genebank (Supplementary Table S1). Seeds of the OzWheat Panel are available for researchers through the AGG.

### Genomic data

Genomic single nucleotide polymorphism (SNP) data was generated from DNA extracted from seedlings for each panel accession. Fresh leaf tissue from seedlings (a pool of 6 plants per line) was freeze-dried and genomic DNA extracted according to Ellis et al. (2005) with the addition of 10µg/ml RNaseA (Sigma, R6513) to the lysis buffer and liquid handling with Microlab NIMBUS robot (Hamilton, Reno, NV, USA). Genotyping with Illumina 90K Infinium iSelect SNP array was performed as outlined in Wang et al. (2014). Alleles were assigned using GenomeStudio (Illumina, San Diego, CA, USA) and a custom Perl script, with SNPs anchored according to sequence alignment with CS Ref Seq v1.0 (IWGSC, 2018). This yielded data for 22,556 polymorphic SNPs across the panel. These were combined with 26,498 SNPs called from transcriptome alignments for the same set of varieties as described by Dillon et al. 2024, to make up a total set of 49,054 SNP markers which were applied in downstream analysis. Transcriptome data used in this study were generated for each panel accession growing under long and short daylength conditions as described by Dillon et al. 2024. Bioinformatic analysis of the transcriptome sequence data produced a matrix of quantitative expression for 44,054 genes across all accessions as described by Dillon et al. 2024, which were applied in downstream analysis.

A SNP Haplotype map (HapMap) file combining 90K and transcriptome SNPs was generated after removal of missing or poor-quality data (genotypes with >50% missing data removed, SNPs with >20% missing data removed) and re-coding to a biallelic score instead of actual nucleotide base (G/C, for HapMap format). Monomorphic markers and those with a minor allele frequency less than 5% were also removed. The HapMap file (sorted by physical chromosome position of SNP in CS Ref Seq v1.0) was used for subsequent association mapping. Graphic visualisation of SNP marker density was produced using the CMPlot package in R (Yin et al., 2021).

### Pedigree and population structure

The Helium Pedigree Visualisation Framework (Shaw et al. 2014) was utilised to view the OzWheat Panel in the context of the wider Australian wheat pedigree. A Helium-compatible text file containing all known ancestors was derived from the International Crop Information System (ICIS) (Portugal et al. 2007) using a custom script for reformatting. Population structure was examined by principal components analysis (PCA) (Patterson et al. 2006) and multidimensional scaling (MDS) in genomics software package TASSEL v5.2.3.1 (Trait Analysis by aSSociation, Evolution and Linkage) (Bradbury et al. 2007). Linkage disequilibrium (LD) was estimated at genome-wide and within -chromosome level in TASSEL using 90K SNPs, filtered to remove unmapped SNPs, those positioned within 10 kbp of each other and those with allele frequency less than 10%. Pairwise associations *(r^2^)* were obtained in a 50 SNP sliding window, excluding heterozygotes. A decay curve was generated by plotting (*r^2^*) against physical distance for a whole-genome representation as well as for each chromosome. To determine the average decay distance per chromosome the background *r^2^* (genome-wide mean) was selected as the threshold of significance, with decay distance being the intercept of this threshold and a fitted decay curve generated in R (locally weighted regression, Loess, R Core Team 2023). These distances were considered when defining the regions of interest from marker-trait associations (MTAs) which were visualised with ChromoMap R package (Anand and Rodriguez, 2022).

### Phenotypic analysis

The OzWheat panel was grown in a polycarbonate greenhouse (CSIRO, Black Mountain) in autumn (shortening days) and spring (lengthening days) of 2016 to best represent local growing conditions, and in long and short days as described by Dillon et al. 2024. In summary, anthesis date (Z61, anthers visible on primary spike) was recorded for each replicate (n=5-6) and to ensure all material was represented in the genome analysis, some winter-types which failed to flower in non-vernalised glasshouse were given a proxy anthesis date (set to be the day after the experiment was harvested).

Field experiments were conducted at CSIRO Ginninderra Experimental Station in Canberra in 2018 and 2019, at Australian Grain Technologies (AGT) Kabinga breeding site, Wagga in 2018 and at the University of Sydney Plant Breeding Institute Narrabri in 2019. Two replicates of each line were sown in a randomised complete block design at each site. In Canberra, each plot comprised 8 rows with 18cm spacing and length of 5 linear metres. Wagga plots comprised 2 rows only, with total plot dimensions 0.75m × 2.5 linear metres and Narrabri configuration was 2m wide, 6-row plots at 3.8m long. Heading date (Z51, date that 50% of plants in the plot had spikes fully emerged from the boot) was recorded, along with plot height at maturity (mean of 3 representative plants per field plot).

### Environment data

Temperature and daylengths for each trial site/year combination were obtained from the SILO Patched Point Dataset, at the nearest stations of the Bureau of Meteorology and Geoscience Australia (Jeffrey et al. 2001, Geoscience Australia 2019). Site descriptions and a summary of the climatic conditions for each site are shown in Supplementary Table S2 and Supplementary Fig. S1. All sites received supplemental irrigation to ensure adequate grain production.

### Statistical analysis

Trait data was analysed, and graphics generated using GenStat version 16.1.0.10916 (VSN International, 2022) and R version 3.2.1 (R Core Team, 2017). To integrate climate data in the field experiments, degree-days to heading (DDTH) was determined by an average equation with a base temperature of 0°C (McMaster and Wilhelm 1997). Raw data from each experiment was checked for normality before fitting a linear mixed model (residual maximum likelihood method, REML) to determine trait values and variance estimates for heritability (Allard, 1999). In the greenhouse, bench and position-within-bench were applied as random factors with genotype as either fixed (to calculate best-linear unbiased estimates (BLUEs)) or random (for predictions (BLUPs)). The analysis of field data included row and column within the block as random effects. To assess the proportion of the genetic to phenotypic variation in the OzWheat panel, and therefore understand the environmental contribution to heading date and height, the genetic and phenotypic coefficients of variation (GCV, PCV) were calculated from variance components (Allard, 1999).

### Association analysis

Genome-wide association analysis (GWAS) was performed in TASSEL v5.2.3.1, with results from different models and correction for population structure compared; a general linear model (GLM) with principal components as covariates in the model (with 1000 permutations), and mixed linear model (MLM) with principal components from MDS analysis, plus kinship matrix based on SNPs included. Quantile-quantile (QQ) and Manhattan plots (CMplot, Yin et al. 2021) were compared for each model in the controlled environment experiments to determine the optimum parameters and significance threshold for genome-wide association analysis of field data. To determine the threshold of significance for associations, Bonferroni correction was used (Kaler and Purcell 2019) comparing significance levels of α (0.05, 0.01, 0.001).

### Data visualisation tool

A standard workflow to visualise the OzWheat SNP and transcriptome data was developed as a Shiny web application (Chang et al. 2024) in the R programming language (R Core Team, 2023) with interactive plot functionality using catmaply (Mauron, 2024). This workflow is illustrated in Supplementary Fig. S2.

## Results

Supplementary data from this study is available at the CSIRO data access portal, https://data.csiro.au/collection/csiro:62968

### OzWheat platform captures significant diversity from the Australian wheat gene pool and global germplasm

Available pedigree information was collated into a Helium-compatible file (Supplementary Table S3) with 1,528 nodes. After filtering, a total of 49,504 SNPs were identified and a kinship matrix for the OzWheat panel derived (Supplementary Table S4, S5). Inclusion of the transcriptome SNPs doubled the marker density achieved by the 90K Illumina array, and included marker saturation in regions which were not well represented by the SNP array alone (for example, close to centromere or on the D-genome, see Supplementary Fig S3).

Principle components analysis (PCA) revealed that the first two components explained 16.9% of the genetic variance (PC1: 11.6%, PC2: 5.8%) with subsequent components explaining below 5% (Supplementary Fig S4). Multi-dimensional scaling (MDS) best resolved tight clusters (Fig. 1) and Scree plot (Supplementary Fig S5) indicated that four components would apply the most stringent conditions to control for population structure in association analysis. Calculation of linkage disequilibrium across the genome indicated an average *r^2^* = 0.24 with decay occurring within 7.3 Mbp (Supplementary Fig. S6, Supplementary Table S6). To interpret subsequent association analysis results, the linkage disequilibrium value for each chromosome defined the regions of interest. That is, for a gene to be considered a candidate from a marker-trait association, it was physically located within the LD estimate for the chromosome identified (see *Association analysis* section below).

**Figure 1.**
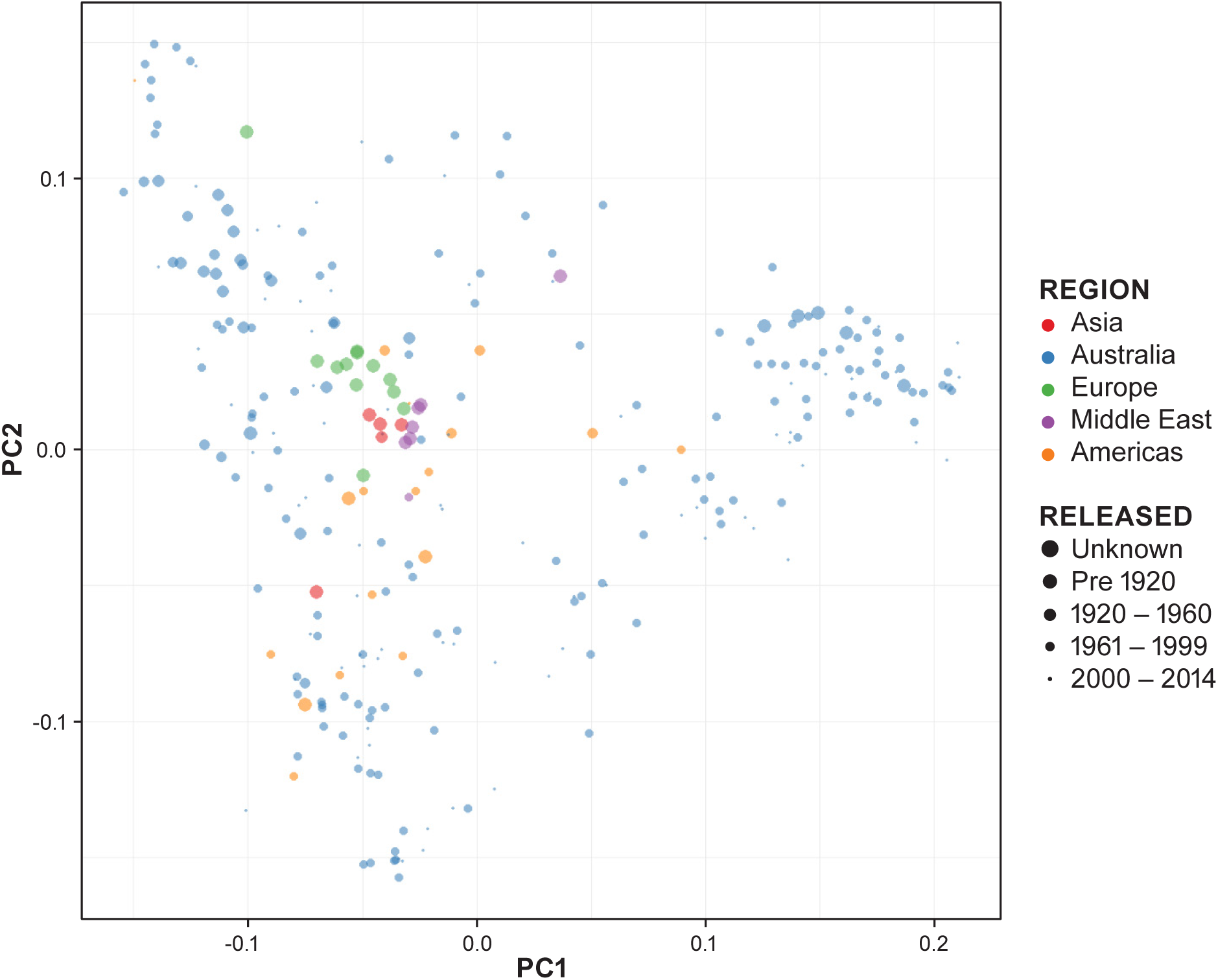
Multi-dimensional scaling (MDS) performed in TASSEL. Principal Co-ordinates Analysis Plot (PCoA) of SNP data in OzWheat, sized by year of release, coloured by region of origin.

### Significant variation for flowering time and plant height in the OzWheat panel

Trait data displayed relatively normal distributions and residual plots revealed that models were appropriate for predictions of days to heading and height and for each trial, and genotype was a highly significant (p<0.001) term in the model (Supplementary Fig S7-10, Supplementary Table S7). Variance components analysis (Table 1-2) within each site revealed that broad-sense heritability was high (from 0.67 to 0.99) and the phenotypic coefficients of variation (PCV) only slightly higher than the genotypic coefficients of variation (GCV), indicating a large effect of genetic background on trait variance. Comparing PCVs for each trait within sites indicated that variability for height was greater than the relative variability for degree-days to heading in Canberra and Wagga, whereas the opposite was true for Narrabri, possibly reflecting the different climatic conditions.

**Table 1.**
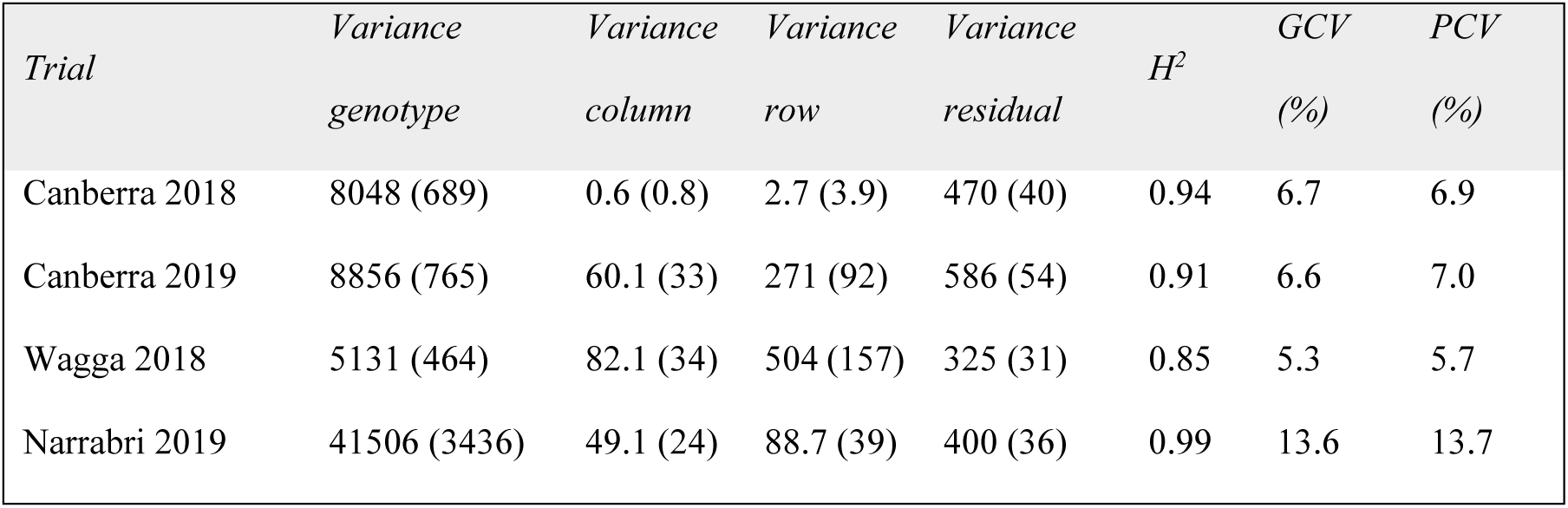
Phenotypic trait analysis, predicted degree-days to heading. Variance components, broad-sense heritability (H^2^), genotypic coefficient of variation (GCV) and phenotypic coefficient of variation (CV) (standard errors in brackets) for OzWheat panel (replicates = 2) grown in Canberra (2018, 2019), Wagga (2018) and Narrabri (2019).

**Table 2.** Phenotypic trait analysis, predicted plot height. Variance components, broad-sense heritability (H^2^), genotypic coefficient of variation (GCV) and phenotypic coefficient of variation (CV) (standard errors in brackets) for OzWheat panel (replicates = 2) grown in Canberra (2018, 2019), Wagga (2018) and Narrabri (2019).

| *Trial* | *Variance*<br>*genotype* | *Variance*<br>*column* | *Variance*<br>*row* | *Variance*<br>*residual* | $H^2$ | *GCV*<br>*(%)* | *PCV*<br>*(%)* |
| --- | --- | --- | --- | --- | --- | --- | --- |
| Canberra 2018 | 98 (9) | 0.003 (0.01) | 0.03 (0.04) | 17 (1.5) | 0.85 | 15.8 | 17.2 |
| Canberra 2019 | 108 (11) | 9 (3.6) | 0.5 (1) | 45 (4.1) | 0.67 | 15.1 | 18.5 |
| Wagga 2018 | 153 (14) | 4 (1.6) | 0.96 (0.73) | 17 (1.6) | 0.88 | 16.3 | 17.4 |
| Narrabri 2019 | 89 (8) | 1.5 (0.7) | 1.4 (0.7) | 11 (1) | 0.86 | 11.9 | 12.9 |

### Association analysis

Different models and significance thresholds for association analysis were compared in controlled environments (Fig. 2). Mixed linear models (MLM) with Bonferroni threshold set at p=0.05, identified 48 marker trait associations (MTAs) (Fig. 3 and listed in Supplementary Tables S8-S9). Manhattan plots for all environments are provided in Supplementary Fig. S11 – 18. A within-environment comparison for flowering time and height GWAS results is shown in Supplementary Fig. S19. Significant MTAs detected in more than one environment (circular Manhattan plot, Fig 4.) were selected for further investigation.

**Figure 2.**
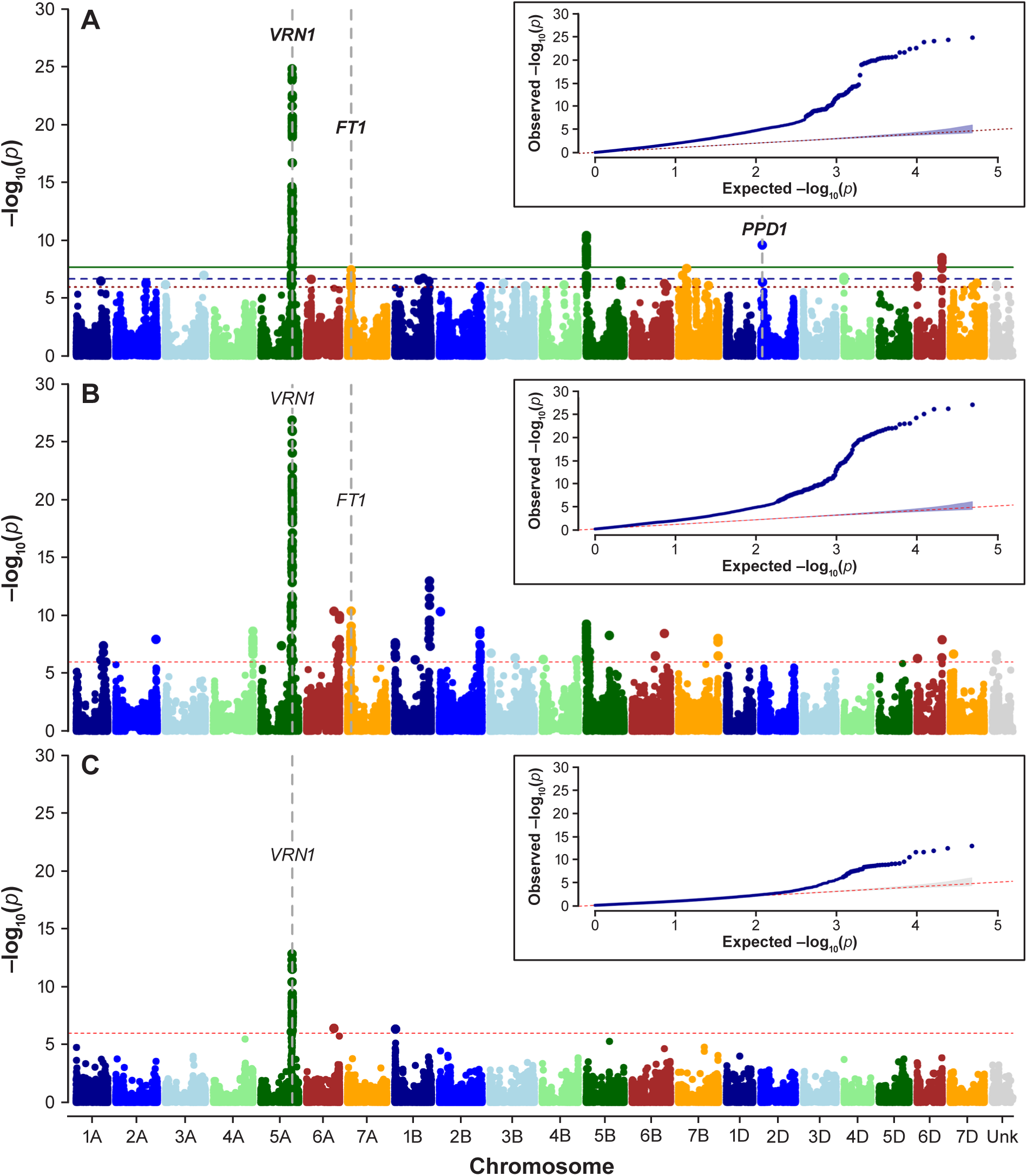
Genome wide association analysis of days to anthesis for plants grown in non-vernalising glasshouse conditions. Location of known major genes for phenology, *VRN1* (chr 5A), *FLOWERING TIME 1 (FT1)* (chr 7A) and *PPD1* (chr 2D) coinciding with MTAs are indicated by grey vertical dashed lines (genome position according to Chinese Spring Ref Seq v1.0). (A). Generalised linear model with 1000 permutations and two principal components for short day experiment (12h). Three levels of significance (Bonferroni thresholds) indicated by red dotted line (0.05), blue dashed line (0.01) and horizontal green line (0.001). (B). Generalised linear model with 1000 permutations and two principal components for long day experiment (16h). Bonferroni level of significance indicated by red dotted line (0.05). (C). Mixed linear model with kinship matrix and four principal components for plants grown in long days (16h). Bonferroni level of significance indicated by red dotted line (0.05).

**Figure 3.**
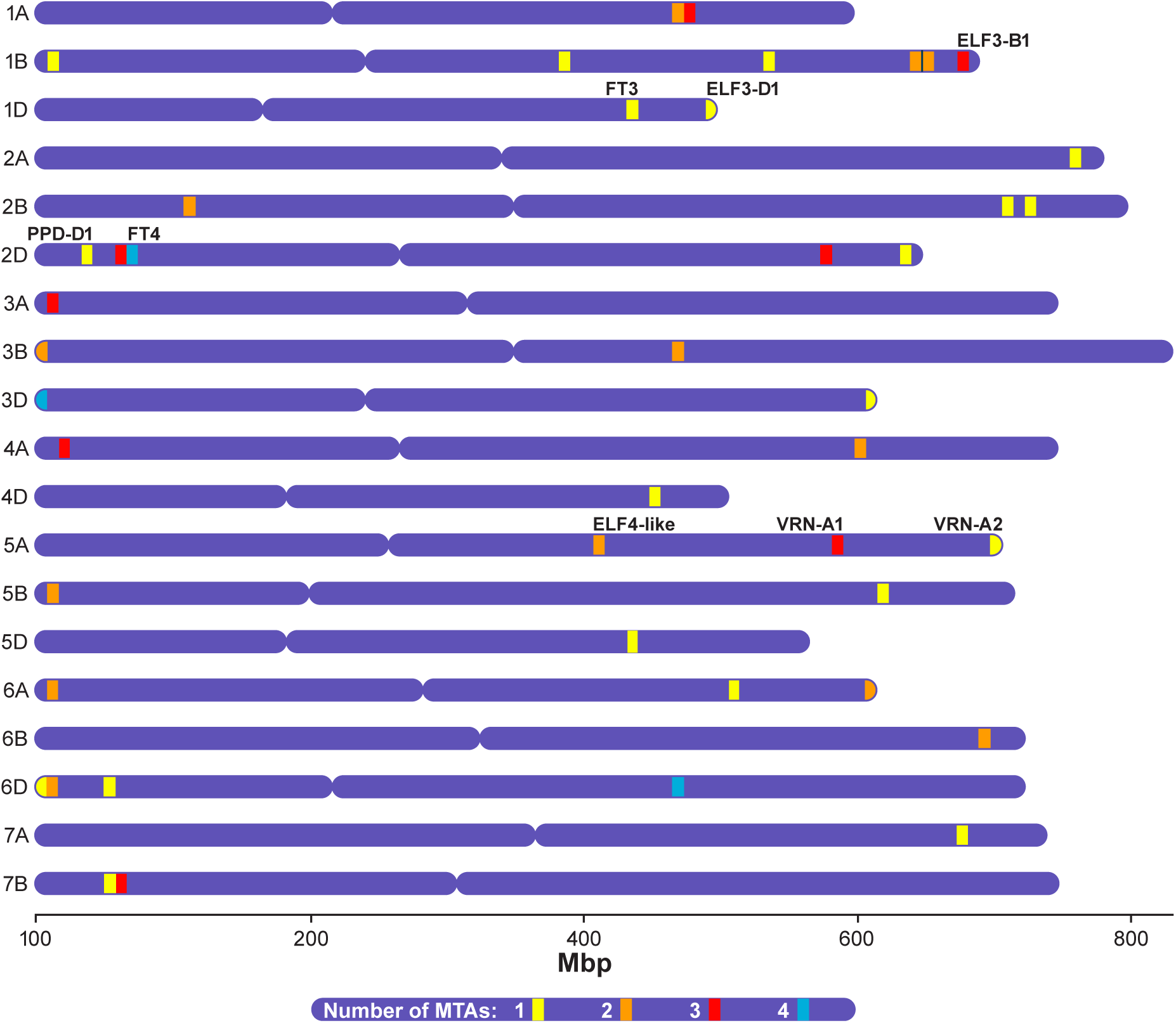
Marker-trait associations for flowering time (mixed linear model, kinship matrix, 4 principal components). Genome position of MTAs identified in controlled conditions (long and short days) and field experiments (Canberra, Wagga, Narrabri 2018 – 2019), relative to known location of major phenological genes in linkage disequilibrium according to physical position in Chinese Spring Ref Seq v1.0.

**Figure 4.**
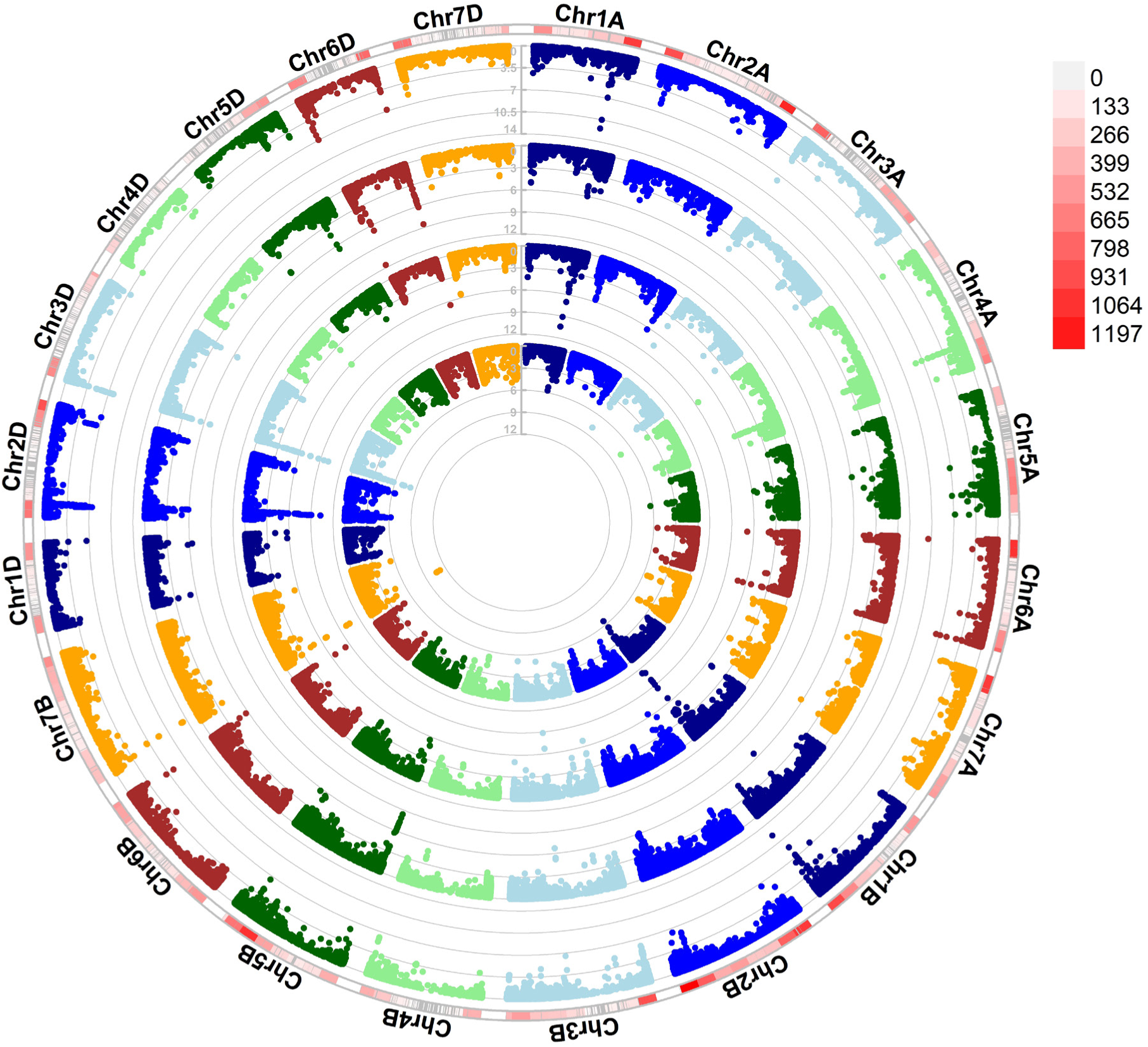
Genome wide association analysis for degree-days to heading at each field location. Circular Manhattan plot (mixed linear model, kinship matrix, 4 principal components); from inner to outer circle, Canberra (2018), Canberra (2019), Wagga (2018), Narrabri (2019). Significance of association (log-10 p-value) designated by grey grid-lines within each site/year, density of SNPs shown in outermost ring (bin size=100MBp) from 0 (white) to 1197 (red).

### Single nucleotide polymorphisms identify candidate genes for adaptation

To identify candidate genes in the OzWheat G2P platform, further evidence aside from marker-trait associations are required. Visualisation of marker alleles flowing through the breeding pedigree can provide confidence that genes associated with adaptation have been identified, since alleles are maintained during the breeding process (Fig. 5A). Alleles of marker SNP2749-1B, associated with flowering time in Canberra and Narrabri (*mDDTH.Cbr19.SNP2749.1B.2* and *mDDTH.Nar19.SNP2749.1B.6*) were present in both winter and spring types, offering potential for this diversity to be utilised in a range of different environments or farming systems. The ability to include gene expression data provides additional support for identification of a candidate gene through GWAS. As shown in Fig. 5B, plants containing contrasting alleles of SNP2749-1B differed in their relative transcript abundance (in crown tissue) when grown in controlled conditions. In addition, we found increased transcript abundance (for both allelic classes) when plants were grown in inductive (long day) conditions relative to short days (data not shown).

**Figure 5.**
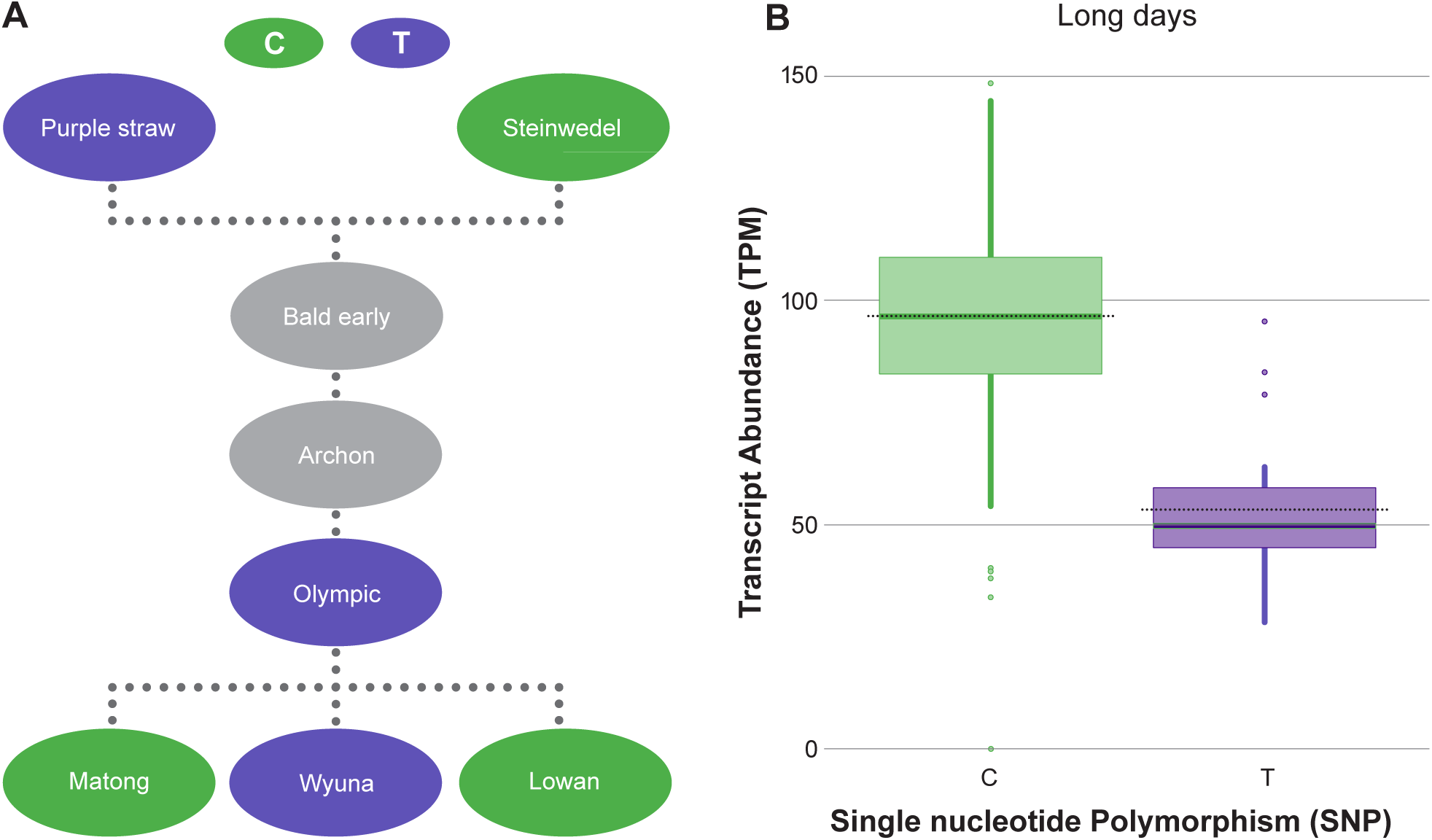
Marker from TraesCS1B01G429200 associated with degree-days to heading at Canberra and Narrabri, SNP2749-1B, alleles in selected OzWheat lines and associated transcript abundance. (A) Green “C” allele and purple “T” allele in historic material and modern cultivars, dashed lines represent simplified crossing schema (not all parents are shown) and SNP data unavailable for grey nodes. (B) Abundance of TraesCS1B01G429200 for the contrasting allelic groups (C/T) in the OzWheat panel, plants grown in long (16h) days.

### Predicted protein sequence

An identified SNP which also encodes an amino acid change or stop codon potentially corresponds to variation that affects function of a gene. The most significant MTA for height in all field environments was identified by SNP21122-4D, located within known dwarfing gene *RHT-D1.* The marker detected a [G/T] point mutation of TraesCS4D01G040400 which induces the premature stop-codon and subsequent truncated protein defined by the *Rht-D1b* dwarfing allele, [T61G] (Peng et al. 1999). We also identified the causal SNP for dwarfing gene *RHT-B1,* (SNP20031-4B, *Rht-B1b*) although this was not associated with plant height in the field.

Aside from *Rht-B1b* and *Rht-D1b* an additional 190 SNPs were identified that induced premature stop codons (nonsense mutations) in the transcriptomes collected from plants grown in controlled conditions, although none of these SNPs were associated with time to flowering or plant height in this study. From 1,3196 missense SNPs (predicted to encode a change in an amino acid) identified in this study, 20 were reported as MTAs for flowering time or height (Supplementary Table S7, S8). For instance, a SNP which encoded an amino acid substitution in TraesCS6D01G028200 was associated with time to flowering at Wagga and the glasshouse. This transcript corresponds to a DExH-box helicase gene, with 80% homology to *BAD REPONSE TO REFRIGERATION 2 (BRR2),* a regulator of flowering time in *Arabidopsis* (Mahrez et al. 2016).

### Coincidence of candidate genes with loci detected in other studies

Transcriptome-derived SNPs are useful to align MTAs and candidate genes identified in other studies. The most significant MTA for flowering time in Canberra and Narrabri was defined by SNPs within TraesCS7B01G055300 (annotated as an ATP-dependent DNA helicase). This transcript was previously reported as a dwarfing gene in wheat (*TaDHL*) through QTL mapping and GWAS (Guo et al. 2022), and additional SNPs were identified in this study (Supplementary Table S10).

### Exploration and visualisation of OzWheat datasets

To explore candidate loci for genome-to-phenome approaches, the Rapid Gene Identification data visualisation tool developed in this study allows users to search and filter the OzWheat database via uploaded lists of SNPs, transcript identifiers, or through a set of dropdown menus. A standard workflow begins by selecting SNPs of interest, for instance those identified through genome-wide association analysis or located in a specific position in the genome (physical position according to Chinese Spring RefSeq v1.0, Alaux et al. 2018). The tool displays SNP information including position, predicted amino acid changes and summary data (allelic calls for the OzWheat panel). The user can explore selected transcripts via a link to the Wheat Expression Browser (Borrill et al. 2016, Ramirez-Gonzalez et al. 2018) and download sequence information to be used for the design of SNP-based markers for example (He et al. 2014). The user can view relative transcript abundances (in short and long days) and allelic diversity within a user-specified window through interactive box plots and heatmaps. With these data visualisations and export functions, the Rapid Gene Identification Tool supports the identification of candidate genes and provides user-friendly access to relevant data which underpins the OzWheat G2P platform.

## Discussion

Functional characterisation of genes in complex polyploid genomes such as wheat is possible through application of high-throughput sequencing technologies and the use of genome-to-phenome (G2P) platforms (Adamski et al, 2020). In this study, use of genetic diversity which is rich in recombinational history provided high-resolution mapping power and identified known genes for adaptation and causal mutations (for instance *Rht-D1b*) in addition to novel loci. The use of important complex traits as the first use-case provided validation of the platform, as well as new biological insights. The most significant region affecting time to flowering when non-vernalised plants were grown in both long and short days coincided with the *VRN1* locus (Fig 2). GWAS using a generalised linear model in these environments also detected SNPs at the region encompassing the *FT1* locus (*VRN3)*, though with less importance relative to *VRN1*. Previous studies identified *FT1* and its interaction with *VRN1* associated with flowering time (Li and Dubcovsky 2008, Deng et al. 2015, DeWitt et al. 2021) and our results support the finding that the A-genome copy, *VRN-A1* has the largest impact on vernalisation requirement compared to the B- and D- genome in Australian wheat (Pugsley 1971, Trevaskis et al. 2003).

We showed the choice of model for association analysis impacted the ability to detect genetic loci. From Fig. 2, the most stringent model (MLM including kinship matrix and 4PCs) produced less-significant marker trait associations and failed to detect the region containing *FT1* associated with time to flowering. It is important therefore, to apply existing knowledge of the genetic architecture of traits if possible. In this case, it is possible that correction for population structure led to the failure to detect *FT1.* Deviation from the 1:1 line of QQ plots as shown in Supplementary Fig. S19 also suggested a difference in significant associations for the different traits (flowering time compared to plant height). It is possible this reflects co-selection of alleles for phenological adaptation. For instance, co-inheritance of non-linked alleles will frequently occur in plants which are well adapted to specific environments due to frequent co-selection of some allelic combinations (for instance, strong vernalisation requirement combined with photoperiod sensitivity to ensure adaptation to environments with cold winters and late frost events). It is also possible that the incorporation of transcript-derived SNPs from tissue that is highly predictive of phenology (the RNA samples included the shoot apical meristem) created a dataset that has a greater proportion of genetic markers associated with flowering time than would be expected by chance. Indeed, all MTAs detected in this study have peak markers derived from the transcriptome rather than the 90K array which suggests some level of bias. This hypothesis will be tested as new transcriptomes from alternate tissues are added to the OzWheat dataset in the future, along with new genotyping information from additional SNP arrays.

In this study a conservative method (Bonferroni correction) was used for thresholding and when comparing different significance levels (α =0.05, 0.01, 0.001) we again found that the ability to detect the region containing *FT1* was lost when levels were greater than 0.01. For this reason, we chose α =0.05 Bonferroni threshold for the field GWAS. The capacity to detect some genes is also limited by alignment of OzWheat transcripts to a single reference (CS RefSeq v1.0). In the future, a *de novo* assembly of the OzWheat pan-transcriptome would overcome a current limitation that only genes which are present in the reference genome are identified.

The use of contrasting controlled environment GWAS is valuable to understand gene by environment interactions and comparisons between plants grown in long and short days identified genes which interact with photoperiod. For instance, known allelic variation at *PPD1* determines if a plant is sensitive or insensitive to the length of days for flowering (Law et al. 1978). Genotypes with daylength sensitivity will be slow to flower, or not flower at all, in short day conditions. We identified the genetic region containing *PPD-D1* in the short day experiment (Fig. 2A), although did not detect its ortholog *PPD-B1.* This suggested the D-genome copy had a greater effect on flowering time in the OzWheat G2P panel (as reported in other studies, see Bentley et al. 2013, Cane et al. 2013). Conversely, when plants were grown in long days, *PPD-D1* was not detected. This is likely due to photoperiod requirement of all plants regardless of their allelic variation being met when grown in this condition (16h days) (Fig. 2B,C).

Another flowering-time MTA (detected on chromosome 5B) was identified when plants were grown in short days in the glasshouse and at Wagga, though the same region was not detected in the long-day experiment (*mDDTH.Wag18.SNP23771.5B.3, mDTA.GHSD.SNP23768.5B.4,* Table S8). This suggested the underlying gene responsible was associated with response to daylength. A cluster of three transcripts are located at this locus in Chinese Spring (*TaBx3B, TaBx4B, TaBx5B*), which are genes involved in synthesis of plant defensive compounds known as benzoxazinones. Genes from this family are also responsive to environmental cues such as daylength and temperature and associated with adaptation (Nomura et al. 2005, Niemeyer 2009, Ben-Abu 2018). A recent transcriptome study revealed that benzoxazinone genes played a role in stem elongation in a mutant with accelerated development, *qd* (Xu et al. 2021) and adaptation to temperate environments during maize domestication (Wang et al. 2017).

From the four field trials conducted in this study, the region containing *VRN1* was only detected as important for time to heading at Narrabri in 2019 (*mDDTH.Nar19.SNP23025.5A.15,* Table S7) which could be explained by the interaction of *VRN1* with temperature. Narrabri recorded the highest minimum temperatures in the field (see Supplementary Fig. S1) which prolonged the time to vernalisation saturation relative to plants grown in Canberra and Wagga. This likely explains the skewed fitted value plot for flowering time residuals (Supplementary Fig. S10B), greatest heritability for degree-days to heading (H^2^=0.99, Table 1) and detection of an MTA linked to *VRN1*.

In field conditions where all vernalisation and photoperiod requirements for the plants are met, variation in time to heading will be due to the effects of *EARLINESS PER SE (EPS)* loci. The identification of such genetic loci is important to consider when fine-tuning adaptation beyond allelic variation for major phenology genes *VRN1* and *PPD1.* The *EPS* gene *EARLY FLOWERING3 (ELF3)* is located at the distal end of group 1 chromosomes (Chinese Spring RefSeq v1.0 A-genome: 591Mbp, B-genome: 685Mbp, D-genome: 493Mbp) and in this study, MTAs for degree days to heading flanked these loci on 1BL (681 – 690Mbp) and 1DL (436 – 495Mbp). Further resolution at these loci is required to determine if *ELF3* underlies the MTAs.

A single transcript identified on chromosome 6AS and orthologous region on 6DS was associated with time to heading at all field sites and the glasshouse (in short days), providing greater confidence that a candidate gene (*BRR2-*like) had been identified. It is possible that the regions on 6A and 6D represent a single locus, since the initial set of SNPs derived from the transcriptome were not filtered for multi-mapped reads. This can lead to hemi-SNPs and subsequently an inability to resolve the genome contribution due to mis-mapped SNPs. Nevertheless, the *BRR2-like* gene is an interesting candidate, a yeast mutant of the RNA helicase *BRR2* was reported to confer cold sensitivity due to a single base-pair substitution within the N-terminal Brr domain (Raghunathan and Guthrie 1998). Mutations in the same domain detected in this study (C-terminal Sec63) were found to affect pre-mRNA splicing through modulation of ATPase activity of the spliceosome (Cordin et al. 2014). In *Arabidopsis, BRR2a* regulated flowering time through disrupted FLOWERING LOCUS C (FLC) splicing (Mahrez et al 2016).

Another helicase gene (ATP-dependent helicase, seed maturation protein TraesCS7B01G055300, 58.7Mbp) was identified for degree-days to heading. This gene was located 30 Mbp distal to *VEGETATIVE TO REPRODUCTIVE TRANSITION 2* (*VRT2*), and more than 270 Mbp from *LATE ELONGATED HYPOCOTYL (LHY)* and *FT1,* so the MTA is unlikely to be associated with these genes known to affect heading date on chromosome 7BS in Chinese Spring. (Yan et al. 2006, Kane et al. 2005, Zhang et al. 2015). Yang et al. (2020) reported a QTL for heading date and yield in a panel of elite Chinese wheat, which maps closeby in Chinese Spring (61.5 Mbp) and it remains to be determined if the region could overlap with a gene associated with flowering in long days, *PPD-B2* reported by Khlestkina et al. (2009). The plant gene expression omnibus database (Koh et al. 2024, available at https://expression.plant.tools/) indicated the helicase transcript in wheat is most highly expressed in the flower bud and coleoptile, and is co-expressed with *VERNALIZATION INSENSITIVE 3* (VIN3, TraesCS1D01G090400) which is associated with chromatin organisation and post-translational histone modification. VIN3 is a polycomb repressive complex (PRC2) induces trimethylation of lysine 27 on histone H3 (H3K27me3) during vernalisation induced flowering of winter cereals (Oliver et al. 2009).

Aside from the association with flowering time in this study, TraesCS7B01G055300 (recently named *TaDHL*) was proposed to influence plant height in wheat (Guo et al. 2022), although we did not detect the region associated with height in the field. Several EMS-derived mutants have been reported at this locus (Krasileva et al. 2017), and future analysis of these lines containing additional SNPs to those already identified might provide further insights into the allelic effects on height and heading date in spring wheat germplasm.

We also found allelic variation which was not explained by winter or spring growth habit. For example, the flow of SNP2749-1B alleles through the breeding pedigree (Fig. 5) suggested no deleterious effects of particular alleles, and that the source of the SNP located in TraesCS1B01G429200 might be Purple Straw. The differences in transcript abundance when the OzWheat population was grouped by this SNP allele suggested lower gene expression for lines carrying the ‘T’ allele compared to ‘C’, with overall gene expression increased in long days (Fig. 5). These results suggested a functional difference between allelic classes, in addition to some interaction with daylength.

The mapping precision of the OzWheat G2P platform was demonstrated when the causal mutation for reduced height was identified (*Rht-D1b*, SNP21122-4D) as the most significant contributor to plant height in all environments (Table S9). We did not find *Rht-B1* associated with plant height in our GWAS, despite the *RhtB1b* allele (identified by SNP20031-4B) present in 46% of lines (compared to 31% of lines in the panel containing *RhtD1b*). A previous GWAS study (Garcia et al. 2019) also did not detect *Rht-B1* as important for height in the field (in southern Australia). We note that the *Rht-B1b* SNP polymorphism [C190T] is also present in the *Rht-B1d* allele derived from Saitama 27 which is prevalent in European wheats and reported to produce taller plants compared to *Rht-B1b* (Pearce et al. 2011, Worland and Petrovic 1988). It is possible that a failure to differentiate *Rht-B1d* and *Rht-B1b* alleles may be confounding our analysis. Additionally, the effects of population structure and other loci or interactions could explain our results. Indeed, Pearce et al. (2011) suggested an alternate mutation outside of the coding region may contribute to height in *Rht-B1d* genotypes, and in the future, additional sequencing or marker screening combined with multi-locus genome wide association analysis might better account for undetected alleles and epistatic effects in the model.

The OzWheat G2P platform has potential to contribute to crop improvement by providing an understanding of genotype by environment interactions via the transcriptome captured in contrasting conditions. This understanding will allow more informed decisions and multiple outputs for breeding. For instance, sequence information provided by the OzWheat G2P platform allows transcript derived markers for breeding to be developed, including kompetitive allele-specific PCR (KASP)s for marker-assisted selection (He et al. 2014; Ramirez-Gonzalez et al. 2015). Additionally, it is possible to identify markers from SNP arrays which are correlated with transcript SNPs and therefore informative for enrichment of favourable alleles during genomic selection (GS). The inclusion of trait-associated transcript markers can improve genomic prediction models for adaptation as demonstrated in maize and rice (Bhandari et al. 2019, Azodi et al, 2019, Wang et al. 2019). An approach which incorporates gene information to improve crop model accuracy is being tested by genetic parameterisation of crop model APSIM (Agricultural Production Systems sIMulator) for improved prediction of wheat phenology (Celestina et al. 2021). Here, the OzWheat G2P platform is being used to predict the cultivar-specific physiological parameters which underpin the model (Dravitzki, 2024 *submitted*) with the aim to provide cross-environment phenology prediction at the time of cultivar release.

To determine the function of candidate genes which have been identified by a G2P platform, investigation of mutants in TILLING (Targeting Induced Local Lesions IN Genomes) populations, analysis of gene expression via transgenics, or gene editing can be deployed (McCallum et al. 2000, Ford et al. 2019). The marker-trait associations identified by the OzWheat G2P platform provide targeted information for sequence capture design to produce new TILLING libraries. Introduction of genetic variation through gene editing for the targets identified in this study is also a path to crop improvement for the outputs of this research. Combined with existing understanding of major genes which contribute to adaptation, there is potential for accelerated genetic gain through multi-targets or “adaptation edits” to be built, which adapt elite cultivars to specific growing environments and markets. For this, an understanding of future climates, farming systems and different end-uses is vital. Examples of some additional traits which could be applied in the OzWheat G2P platform and targeted for a gene editing package include plant architecture (for example, short-stature wheat with a long coleoptile), water-use efficiency, tolerance to temperature extremes, improved grain quality and disease resistance. That, which would traditionally take many years and an entire breeding program to deliver, could now be more achievable through gene editing for crop improvement, and ultimately, comparative genomics linking multiple G2P platforms in different species to produce a crop-agnostic system might even be possible.

## Conclusion

This study delivered the OzWheat genome-to-phenome (G2P) platform for wheat pre-breeding and research which is accessible through germplasm, data and visualisation tools. We demonstrated the power of genome-wide association studies in contrasting controlled conditions and multiple environment field trials to detect novel loci which underpin adaptive traits, and to understand genotype by environment interactions. A high degree of mapping resolution was achieved, and since the OzWheat panel was curated to capture breeding history, the loci detected were relevant in a broad range of genetic diversity. In addition to the sequence variation captured, gene expression information from the transcriptome provided a powerful tool for functional genomics, and candidate genes identified in this pilot study have potential to contribute to the development of adapted wheat suitable for changing global climates. We propose the OzWheat G2P platform is a re-usable and expandable resource for the wheat research community. Additional trait and ‘omics data layers will meet new science challenges and answer different biological questions in the future, and as the dataset expands, new methods for integration and analysis provide further insight into the genetic basis of adaptation and other traits.

## Acknowledgments

This research was funded by CSIRO Agriculture & Food, CSIRO IM&T Scientific Computing, and the Australian Government Research Training Program (RTP) in collaboration with University of Sydney (J. Hyles PhD project, Genome-to-Phenome: Diversity for Adaptation and Phenology in Wheat), Australian Grain Technologies (AGT) and the Grains Research & Development Corporation (CSP00183 Pedigree-based Association Analysis of Wheat Phenology). The authors would like to acknowledge the contribution of Dr Paul Shaw (James Hutton Institute) and Mr Septian Razi (CSIRO Data61) for assistance with pedigree visualisation, Mr Carl Davies for graphic design and biostatistician Dr Alec Zwart for analytics expertise. We also thank Dr Sally Norton and Dr Brett Lobsey (Winter Cereals Collection and Australian Grains Genebank) for curation assistance. We acknowledge the data collection from field trial teams lead by Mr Graeme Rapp (University of Sydney Plant Breeding Institute, Narrabri), Mr Brett Irons (AGT) and Mr Tom McLucas (Ginninderra Experimental Station, CSIRO, Canberra). Finally, we thank the generations of breeders, researchers, and farmers who developed, selected and collected the wheat genetics used in this study.

